# Genome-wide association mapping and genomic prediction unravels CBSD resistance in a *Manihot esculenta* breeding population

**DOI:** 10.1101/158543

**Authors:** Siraj Ismail Kayondo, Dunia Pino Del Carpio, Roberto Lozano, Alfred Ozimati, Marnin Wolfe, Yona Baguma, Vernon Gracen, Offei Samuel, Morag Ferguson, Robert Kawuki, Jean-Luc Jannink

## Abstract

Cassava *(Manihot esculenta* Crantz), a key carbohydrate dietary source for millions of people in Africa, faces severe yield loses due to two viral diseases: cassava brown streak disease (CBSD) and cassava mosaic disease (CMD). The completion of the cassava genome sequence and the whole genome marker profiling of clones from African breeding programs (www.nextgencassava.org) provides cassava breeders the opportunity to deploy additional breeding strategies and develop superior varieties with both farmer and industry preferred traits. Here the identification of genomic segments associated with resistance to CBSD foliar symptoms and root necrosis as measured in two breeding panels at different growth stages and locations is reported. Using genome-wide association mapping and genomic prediction models we describe the genetic architecture for CBSD severity and identify loci strongly associated on chromosomes 4 and 11. Moreover, the significantly associated region on chromosome 4 colocalises with a *Manihot glaziovii* introgression segment and the significant SNP markers on chromosome 11 are situated within a cluster of nucleotide-binding site leucine-rich repeat (NBS-LRR) genes previously described in cassava. Overall, predictive accuracy values found in this study varied between CBSD severity traits and across GS models with Random Forest and RKHS showing the highest predictive accuracies for foliar and root CBSD severity scores.

## INTRODUCTION

Cassava *(Manihot esculenta* Crantz), is a major source of income and dietary calories for more than 800 million people across the globe especially in Sub Saharan Africa (SSA) and recently, due to the unique starch qualities of the storage roots cassava is also turning into an industrial crop (Pérez *et al.,* 2011). Although cassava is a resilient crop, its production is threatened by viral diseases such as Cassava brown streak virus disease (CBSD), which causes major yield losses to poor farming families (ASARECA:, 2013; Ndunguru *et al.,* 2015; Patil *et al.,* 2015). CBSD is caused by two major strains; *Cassava brown streak virus* (CBSV) and *Ugandan cassava brown streak virus* (UCBSV) both CBSVs have successfully colonized the lowland and highland altitudes across East Africa and new strains are emerging (Winter *et al.,* 2010; Ndunguru *et al.,* 2015; Alicai *et al.,* 2016). In Uganda, because of CBSVs and agronomical practices, cassava yields were recorded to be eight times lower than the yield potential for this crop (ASARECA:, 2013).

In addition to the uncontrolled exchange of infected cassava stakes among farmers across borders, CBSVs are transmitted by the African whitefly (*Besimia tobaci)* in a semi-persistent manner (Legg, Sseruwagi, *et al.*, 2014; McQuaid *et al.*, 2016). Upon infection, the viruses use the transport system of the plant and cause yellow chlorotic vein patterns along minor veins of leaves in susceptible cassava clones (Ogwok *et al.*, 2010; Maruthi *et al.*, 2016; Anjanappa *et al.*, 2016). On the stem, prominent brown elongated lesions commonly referred to as “brown streaks” are formed and in the storage roots, necrotic hard-corky layers are formed in the root cortex of the most susceptible cassava clones (Hillocks *et al.*, 1996; Legg, Somado, *et al.*, 2014; Ndyetabula *et al.*, 2016).

Earlier, CBSD resistance breeding initiatives have highlighted the polygenic nature of inheritance in both intraspecific and interspecific cassava hybrids (Nichols, 1947; Hillocks and Jennings, 2003; Munga, 2008; Kulembeka, 2010). In view of the rapid virus evolution and the insufficiency of dependable virus diagnostic tools (Alicai *et al.*, 2016) breeding for durable CBSD resistance, has been the main strategy to control CBSD spread in Eastern Africa. Most of the available elite cassava lines have exhibited some level of sensitivity to CBSVs ranging from mild sensitivity to total susceptibility. Moreover, clones classified as resistant and tolerant show diverse symptom expression, restricted virus accumulation or recovery after clonal propagation (Hillocks and Jennings, 2003; Alicai *et al.*, 2016).

Overall, in cassava for many traits the rate of genetic improvement following a traditional breeding pipeline has been slower due to the combination of several biology-related issues such as: poor flowering, length of breeding cycle, limited genetic diversity and slow rate of multiplication of planting materials.

Recently, using genotypic and phenotypic information genome wide association mapping (GWAS) has been used to unravel the genetic architecture of cassava mosaic disease (CMD) (Wolfe *et al.*, 2016) and beta carotene content (Esuma *et al.*, 2016). Both studies have been successful in identifying associated loci with traits of interest. In addition, the performance of genomic prediction for different traits was previously evaluated using historical phenotypic and genotyping by sequencing (GBS) datasets from the International Institute of Tropical Agriculture in Nigeria (Elshire *et al.*, 2011; Ly, Hamblin, Rabbi, Melaku, Bakare, Gauch, *et al.*, 2013). Genomic Selection (GS) is a breeding method alternative to marker assisted selection and conventional phenotypic selection which can accelerate genetic gains through the use of phenotypic and genotypic data from a training population (Meuwissen *et al.*, 2001; Jannink *et al.*, 2010; Lorenz *et al.*, 2011). The performance of different GS models has been evaluated in various species and in many traits (Resende *et al.*, 2012; Gouy *et al.*, 2013; Heslot *et al.*, 2014; Charmet *et al.*, 2014; Cros *et al.*, 2015). Recently the potential of GS for CMD resistance has been reported with predictive accuracies ranging from 0.53 to 0.58 (Wolfe *et al.*, 2016).

In the present study we followed a GWAS approach in combination with genomic prediction to unravel the genetic architecture of CBSD in two Ugandan breeding populations. While one of our main objectives was to assess the current predictive accuracy for CBSD we also aimed to identify the most promising genomic prediction models that can account for CBSD genetic architecture. GWAS identified loci strongly associated with CBSVs resistance to foliar symptoms which co-locate with an introgression block from a cassava wild progenitor, *M. glaziovii* (Bredeson *et al.*, 2016) and with root necrosis which were close to a cluster of plant defence response-related genes annotated in the cassava genome (Lozano *et al.*, 2015). The presence of introgressions segments from the wild progenitors into the elite breeding lines is the result of cassava improvement programs at the Amani Research Station throughout the 1940s and 1950s (Jennings and Iglesias, 2002; Hillocks and Jennings, 2003).

Here we demonstrated with the synergistic implementation of GWAS and GS that GWAS could be used as a prioritization tool to identify markers for genomic prediction for CBSD resistance in cassava. In addition to unravelling the genetics of CBSD resistance these findings may help in the identification of significant causal polymorphisms to guide marker-assisted breeding for CBSD severity that may greatly improve cassava breeding in the face of increasing disease threats to agricultural production.

## MATERIALS AND METHODS

### Plant material

Phenotypic data was collected from two GWAS panels (Supplementary table 1), GWAS panel 1 composed of 429 clones and GWAS panel 2 which was composed of 872 clones. The combined dataset of 1281 cassava clones were developed through three cycles of genetic recombination between cassava introductions and local elite lines by the National root crops breeding program at NaCRRI. These cassava clones have a diverse genetic background whose pedigree could be traced back to introductions from the International Institute of Tropical Agriculture (IITA), International Center for Tropical Agriculture (CIAT) and the Tanzania national cassava breeding program (Supplementary table 1).

### Phenotyping

The GWAS panel trials were conducted in five locations; Namulonge, Kamuli, Serere, Ngetta and Kasese in Uganda.

GWAS panel 1 data was collected in two years across three locations, each trial was designed and laid out as a 6 by 30 alpha-lattice design with two-row plots of five plants each at a spacing of 1 meter by 1 meter. GWAS panel 2 was evaluated in three locations, on each location, five rows of test clones were bordered by two CBSD susceptible clones in order to increase CBSD disease pressure (TME204). Clones from GWAS panel 2 were evaluated as single entries per location being connected by six common checks in an augmented completely randomized block design with 38 blocks per site (Federer *et al.*, 2002; Federer and Crossa, 2012).

CBSD severity was scored at 3 (CBSD3S), 6 (CBSD6S), and 9 (CBSD9S) months after planting (MAP) for foliar and 12 MAP (CBSDRS) for root symptoms respectively. The CBSD9S scores were not available for GWAS panel 1.

CBSD severity was measured based on a 5-point scale with a score of 1 implying asymptomatic conditions and a score 5 implying over 50% leaf vein clearing under foliar symptoms. However, at 12 MAP a score of 5 implies over 50% of root-core being covered by a necrotic corky layer. (Supplementary Figure 1)

Clones were classified with a score of 5 if pronounced vein clearing at major leaf veins were jointly displayed with brown streaks on the stems and shoot die-back that appeared as a candle-stick. Clones with 31 – 40% leaf vein clearing together with brown steaks at the stems were classified under score 4. A Score of 3 was assigned to clones with 21 – 30% leaf vein clearing with emerging brown streaks on the stems. While a score of 2 was assigned to clones that only displayed 1 – 20% leaf vein clearing without any visible brown streak symptoms on the stems. Plants classified with a score of 1 showed no visible sign of leaf necrosis and brown streaks on the stems. On the other hand, root symptoms were also classified into 5 different categories based on a 5 – point standard scale (Jennings and Iglesias, 2002; Hillocks and Jennings, 2003).

### Two-stage genomic analyses

For the two stage analyses, the first stage involved accounting for trial-design using a linear mixed model to obtained de-regressed BLUPs (drgBLUPs) and the second stage involved the use of de-regressed BLUPs in GWAS and Genomic prediction.

For the panel 1 we fitted the model: =**X***β* + **Z_clone_***c* + **Z_range(loc)_***r* + **Z_block(range)_***b* + ε, using the *lmer* function from the *lme4* R package (Bates et al., 2015). In this model, ***β*** included a fixed effect for the population mean and location. The incidence matrix **Z_clone_** and the vector *c* represent a random effect for clone c∼N(0, 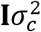) and **I** represent the identity matrix. The range variable, which is the row or column along which plots are arrayed, is nested in location-rep and is represented by the incidence matrix **Z_range(loc.)_** and random effects vector r∼N(0, 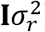). Block effects were nested in ranges and incorporated as random with incidence matrix Z_block(range)_ and effects vector *b*∼N(0,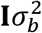). Residuals *ε* were fit as random, with ε∼N(0, 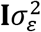).

For panel 2 we fitted the model ***y*** = **X**β + **Z_clone_***c* + **Z_block_***b* + *ε* Where *y* was the vector of raw phenotypes, ***β*** included a fixed effect for the population mean and location with checks included as a covariate. The incidence matrix **Zclone** and the vector *c* are the same as the aforementioned model and the blocks were also modeled with incidence matrix **Z_block_** and **b** represents the random effect for block. The best linear predictors (BLUPs) of the clone effect (ĉ) were extracted as de-regressed BLUPS following the formula (Garrick *et al.*, 2009):

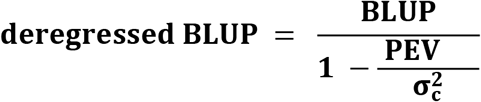

Where PEV is the prediction error variance of the BLUP and 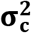is the clonal variance component.

### DNA preparation and Genotyping by sequencing (GBS)

Total genomic DNA was extracted from young tender leaves of all cassava clones included in the phenotyping trials according to standard procedures using the DNAeasy plant mini extraction kit (QIAGEN, 2012). Genotyping-by-sequencing (GBS)(Elshire *et al.*, 2011) libraries were constructed using the ApeKI restriction enzyme (Hamblin and Rabbi, 2014). Marker genotypes were called using TASSEL GBS pipeline V4 (Glaubitz *et al.*, 2014) after aligning the reads to the Cassava v6 reference genome (Prochnik *et al.*, 2012; Goodstein *et al.*, 2014). Variant Calling Format (VCF) files were generated for each chromosome. Markers with more than 60% missing calls were removed. Genotypes with less than five reads were masked before imputation. Additionally, only biallelic SNP markers were considered for further processing.

The marker dataset consisted of a total of 173,647 bi-allelic SNP markers called for 986 individuals. This initial dataset was imputed using Beagle 4.1 (Browning and Browning, 2016). After timputation, 63,016 SNPs had an AR2 (Estimated Allelic r-squared) higher than 0.3 and were kept for analysis; from these, 41,530 had a minor allele frequency (MAF) higher than 0.01 in our population. Dosage files for this final dataset were generated and used for both GWAS and GS analyses.

### Genetic correlations and heritability estimates

Correlation across CBSD traits was estimated using pairwise correlations for each location using the drgBLUPs values obtained after fitting the aforementioned linear mixed model. Broad sense heritabilities (plot-mean basis) were calculated using the estimated variance components from the first step of the two-step genomic analysis as explained previously.

In addition,SNP-based heritabilities were calculated for each GWAS panel by fitting a single-step mixed-effects model, the full models which specified clone as a random effect were fitted using the *emmreml* function from the *EMMREML* R package (Akdemir and Okeke, 2015). The random effect was modeled as having co-variance proportional to the kinship matrix, which was calculated using the *A.mat* function from the *rrBLUP* R package (Endelman, 2011).

### Genome-wide association analysis for CBSV severity

Although pedigree records indicate the two GWAS panels to be closely related a principal component analysis (PCA) was performed in order to characterize these panels and to identify any population stratification between the two GWAS panels. We used the imputed dataset of 63,016 SNP markers to calculate the PCs with the function *princomp* in R.

With the imputed dataset of 63,016 SNP markers and 986 individuals genome wide association was performed using a mixed linear model association analysis (MLMA) accounting for kinship as implemented in GCTA (v 1.26.0) (Yang *et al.*, 2011). Specifically, we followed a leave one chromosome out approach, with this approach the chromosome on which the candidate SNP markers are tested gets excluded from the genomic relationship (GRM) calculation. Bonferroni correction (reference) was used to correct for multiple testing with a significance threshold set at 5.9. Manhattan plots with transformed -log_10_(P-value) were generated using R package *qqman* (Turner, 2014).

### Genomic prediction models

To assess the potential of implementing genomic selection for CBSD, seven genomic prediction models were keenly examined; genomic best linear unbiased prediction (GBLUP), reproducing kernel Hilbert spaces (RKHS), BayesCpi, Bayesian LASSO, BayesA, BayesB and Random forest (RF).

**GBLUP**. In this prediction model, the GEBVs are obtained after fitting a linear mixed model where the genomic realized relationship matrix is based on SNP marker dosages. Accordingly, the genomic relationship matrix was constructed using the function *A.mat* in the R package rrBLUP (Endelman, 2011) and follows the formula of VanRaden (2008), method two. GBLUP predictions were made with the function *emmreml* in the R package EMMREML (Akdemir and Okeke, 2015).

**Multi-kernel GBLUP**. Because the most significant QTLs for foliar severity 3 and 6 MAP were mapped on chromosomes 4 and 11 (this paper) we followed a multi-kernel approach by fitting three kernels with genomic relationship matrices constructed with SNP markers from chromosomes 4 (G_chr4_), 11 (G_chr11_) and SNPs from the other chromosomes (G_allchr-[4,11]_). Multikernel GBLUP predictions were made with the function *emmremlMultiKernel* in the R package EMMREML (Akdemir and Okeke, 2015).

**RKHS**. Unlike GBLUP for RKHS we use a Gaussian kernel function: *Kij* = exp (-(d_ij_ θ)), where *K_ij_* is the measured relationship between two individuals, *d_ij_* is their Euclidean genetic distance based on marker dosages and θ is a tuning (“bandwidth”) parameter that determines the rate of decay of correlation among individuals. This function is nonlinear and therefore the kernels used for RKHS can capture non-additive as well as additive genetic variation. To fit a multiple-kernel model with six covariance matrices we used the *emmremlMultiKernel* function in the EMMREML package, with the following bandwidth parameters: 0.0000005, 0.00005, 0.0005, 0.005, 0.01, 0.05 (Multi-kernel RKHS) and allowed REML to find optimal weights for each kernel.

**Bayesian maker regressions**. We tested four Bayesian prediction models: BayesCpi (Habier *et al.*, 2011), the Bayesian LASSO (Park and Casella, 2008), BayesA, and BayesB (Meuwissen *et al.*, 2001). The Bayesian models we tested allow for alternative genetic architectures by way of differential shrinkage of marker effects. We performed Bayesian predictions with the R package BGLR (Pérez and De Los Campos, 2014)

**Random Forest**. Random forest (RF) is a machine learning method used for regression and classification (Breiman, 2001; Strobl *et al.*, 2009; Charmet and Storlie, 2012). Random forest regression with marker data has been shown to capture epistatic effects and has been successfully used for prediction (Breiman, 2001; Motsinger-Reif *et al.*, 2008; Heslot *et al.*, 2012; Charmet *et al.*, 2014; Spindel *et al.*, 2015). We implemented RF using the random Forest package in R (Liaw and Wiener, 2002) with the parameter, *ntree* set to 500 and the number of variables sampled at each split *(mtry)* equal to 300.

### Introgression Segment Detection

To identify the genome segments in the two GWAS panels, we followed the approach described in Bredeson et al. (Bredeson *et al.*, 2016). We used the *M. glaziovii* diagnostic markers identified in Supplementary Dataset 2 of Bredeson et al. (Bredeson *et al.*, 2016), these ancestry diagnostic (AI) SNPs were identified as being fixed for different alleles in a sample of two pure *M. esculenta* (Albert and CM33064) and two pure *M. glaziovii.*

Out of 173,647 SNP in our imputed dataset, 12,502 matched published AI SNPs. For these AI SNPs, we divided each chromosome into non-overlapping windows of 20 SNP. Within each window, for each individual, we calculated the proportion of genotypes that were homozygous (G/G) or heterozygous (G/E) for *M. glaziovii* allele and the proportion that were homozygous for the *M. esculenta* allele (E/E). We assigned G/G, G/E or E/E ancestry to each window, for each individual only when the proportion of the most common genotype in that window was at least twice the proportion of the second most common genotype. We assigned windows a “No Call” status otherwise.

We also used this approach on six whole-genome sequenced samples from the cassava HapMap II (Ramu *et al.*, 2016). These included the two “pure cassava” and *M. glaziovii* (S) from Bredeson et al. (Bredeson *et al.*, 2016), plus an additional *M. glaziovii*, and two samples labeled Namikonga. Because these samples came from a different source from most our samples, we could find only 11,686 SNPs that matched both the sites in the rest of our study sample and the list of ancestry informative sites for analysis.

### Linkage disequilibrium plots

To confirm whether a large haplotype block present on chromosome 4 colocate with a GWAS QTL identified on this chromosome we calculated LD scores of every SNP marker on chromosome 4 in a 1Mb window using GCTA (Yang *et al.*, 2011). Briefly, LD score for a given marker is calculated as the sum of R^2^ adjusted between the index marker and all markers within a specified window. The adjusted R^2^ is an unbiased measure of LD:

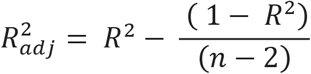

Where “n” is the population size and R^2^ is the usual estimator of the squared Pearson’s correlation (Bulik-Sullivan *et al.*, 2015). The resulting LD scores were then plotted against the GWAS log_10_ (Pvalue) of every marker on chromosome 4.

To highlight the importance of the associated markers on chromosome 11 we calculated pairwise squared Pearson’s correlation coefficient (r^2^) between the top significant GWAS SNP hit on this chromosome and neighboring markers in a window of 2Mb (1Mb upstream and 1Mb downstream). (plink ref)

### Candidate gene identification

We used the mlma GCTA output to filter out SNP markers based on -log10 (P-value) values higher than the Bonferroni threshold (∼ 5.9). The resulting significant SNP markers were then mapped onto genes using the SNP location and gene description from the M.esculenta_305_v6.1.gene.gff3 available in Phytozome 11(Goodstein et al., 2014) for *Manihot esculenta* v6.1 using the intersect function from bedtools (Quinlan and Hall, 2010).

## RESULTS

### Phenotypic variability for severity to cassava brown streak virus infection

In the present study field disease scoring was done based on a standard CBSD scoring scale that ranges from 1 to 5 for both foliar and root symptoms (Supplementary figure 1).

Datasets for CBSD foliar and root severities of the evaluated germplasm are presented in Supplementary figures 2 and 3, both GWAS panels exhibited differential response to CBSVs at three,six, nine and twelve months as revealed in the great variability of the deregressed BLUPs. Interestingly, clones which displayed an intermediate response were by far more abundant than clones with susceptible or resistance response.

Phenotypic correlations for foliar and root severities (CBSD3S, CBSD6S and CBSDRS) within panels and within and across locations are presented in Supplementary figure 4, Supplementary tables 2 and 3 with clear differences in CBSD severity scores.

For panel 1, results varied across locations and CBSD severity traits the lowest correlation value was between Ngetta and Kasese (0.09) and the highest between Namulonge and Kasese (0.60) both values correspond root severity scoring (Supplementary table 2A).

For panel 2 the results varied across locations and CBSD severity traits with correlation values ranging between -0.08 for CBSD9S (Namulonge-Kamuli) and 0.51 CBSD3S (Kamuli-Serere) (Supplementary table 2B).

Within locations across traits the highest correlation values were found in panel 1 for foliar scorings CBSD3S and CBSD6S (r^2^ > 0.5) (Supplementary table 3A). For panel 2, correlation across traits varied depending on the location, nonetheless correlations across foliar traits were generally higher than those between foliar and root severity (Supplementary table 3B).

Heritability estimate values for CBSD3S, CBSD6S and CBDRS were low to intermediate with broad-sense heritability (H^2^) estimates spanning a wide range (11% to 73%) for both panels across locations (Table 1). For GWAS panel 2, broad-sense heritability (H^2^) estimates ranged between 56% and 63% for CBSD3S and between 60% and 62% for CBSD6S; while for GWAS panel 1 ranged between 11% and 51%.

**Table 1.**
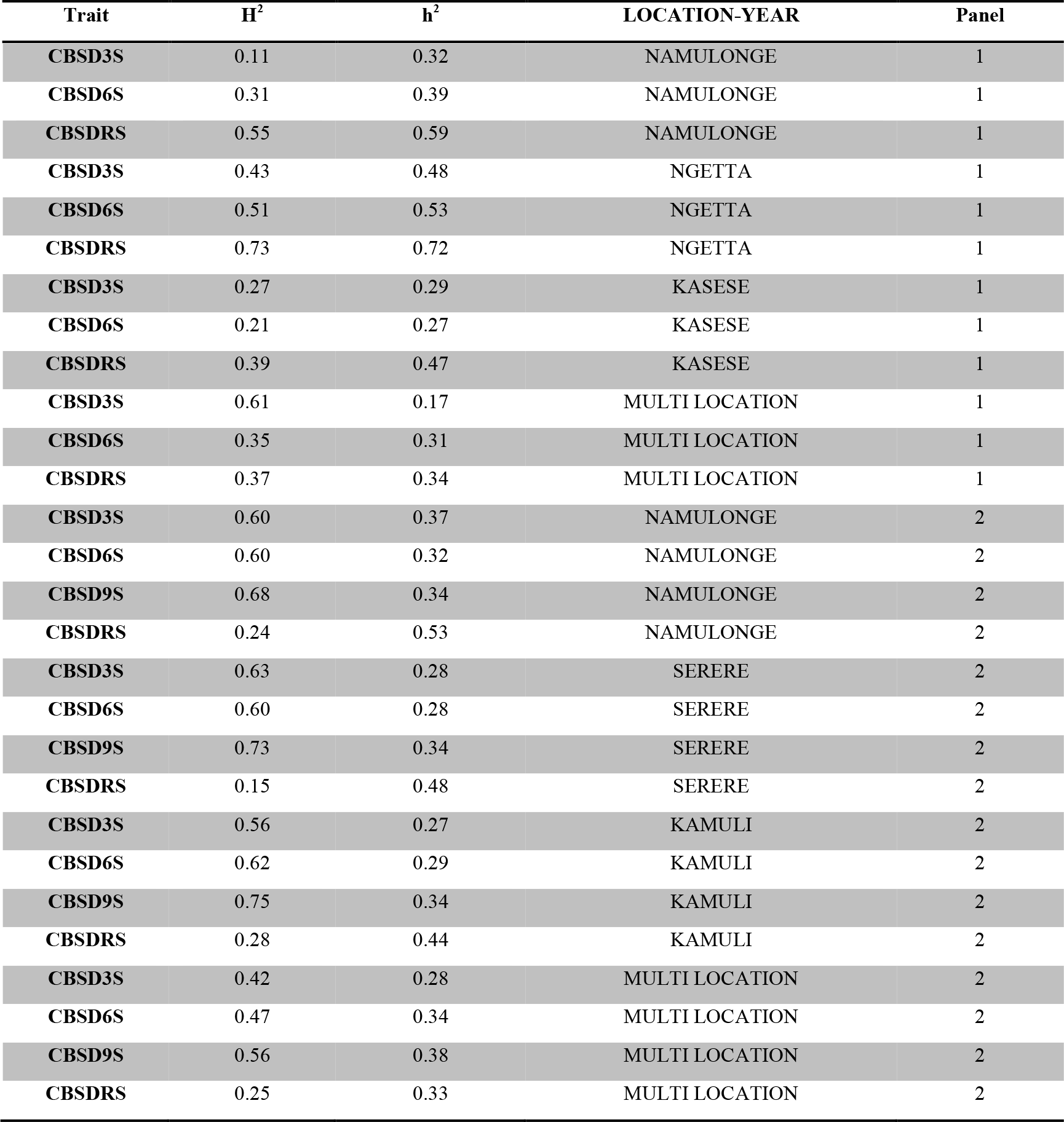
Broad sense heritability (H^2^) and SNP heritability (h^2^) of foliar and root CBSD severity. Broad-sense heritability (H^2^) values were calculated using the variance components obtained from a model fitted using the *Imer* function from the lme4 R package. SNP heritability values were calculated using the variance components obtained obtained from a model fitted using the EMMREML R package. Heritability values estimates were calculated for sets 1 and 2 separately.

| Trait | $H^2$ | $h^2$ | LOCATION-YEAR | Panel |
| --- | --- | --- | --- | --- |
| CBSD3S | 0.11 | 0.32 | NAMULONGE | 1 |
| CBSD6S | 0.31 | 0.39 | NAMULONGE | 1 |
| CBSDRS | 0.55 | 0.59 | NAMULONGE | 1 |
| CBSD3S | 0.43 | 0.48 | NGETTA | 1 |
| CBSD6S | 0.51 | 0.53 | NGETTA | 1 |
| CBSDRS | 0.73 | 0.72 | NGETTA | 1 |
| CBSD3S | 0.27 | 0.29 | KASESE | 1 |
| CBSD6S | 0.21 | 0.27 | KASESE | 1 |
| CBSDRS | 0.39 | 0.47 | KASESE | 1 |
| CBSD3S | 0.61 | 0.17 | MULTI LOCATION | 1 |
| CBSD6S | 0.35 | 0.31 | MULTI LOCATION | 1 |
| CBSDRS | 0.37 | 0.34 | MULTI LOCATION | 1 |
| CBSD3S | 0.60 | 0.37 | NAMULONGE | 2 |
| CBSD6S | 0.60 | 0.32 | NAMULONGE | 2 |
| CBSD9S | 0.68 | 0.34 | NAMULONGE | 2 |
| CBSDRS | 0.24 | 0.53 | NAMULONGE | 2 |
| CBSD3S | 0.63 | 0.28 | SERERE | 2 |
| CBSD6S | 0.60 | 0.28 | SERERE | 2 |
| CBSD9S | 0.73 | 0.34 | SERERE | 2 |
| CBSDRS | 0.15 | 0.48 | SERERE | 2 |
| CBSD3S | 0.56 | 0.27 | KAMULI | 2 |
| CBSD6S | 0.62 | 0.29 | KAMULI | 2 |
| CBSD9S | 0.75 | 0.34 | KAMULI | 2 |
| CBSDRS | 0.28 | 0.44 | KAMULI | 2 |
| CBSD3S | 0.42 | 0.28 | MULTI LOCATION | 2 |
| CBSD6S | 0.47 | 0.34 | MULTI LOCATION | 2 |
| CBSD9S | 0.56 | 0.38 | MULTI LOCATION | 2 |
| CBSDRS | 0.25 | 0.33 | MULTI LOCATION | 2 |

Narrow-sense heritability (h^2^), also referred to as SNP heritability, was estimated using the variance components obtained as a result of fitting a one step model using the genetic relationship matrix (GRM) for each panel. For panel 1, the broad-and narrow-sense heritability values were comparable across locations except for the multi-location model. For panel 2, for most locations the broad-sense heritability estimates were larger than the narrow-sense heritability estimates. The high variability observed within and across GWAS panels reflects differences in population composition, field design and environmental effects.

### Genome wide association mapping for CBSV severity in cassava

The extent of subpopulation structure between the two GWAS panels was examined by PCA, which showed no distinct clusters: clones from both panels had mixed distribution. Overall, the first three PCAs accounted for 60% of the genetic variation observed in the data (Figure 1). The first PC accounted for 30% of the observed variation while the second and third PCs contributed 20% and 10% respectively.

**Figure 1.**
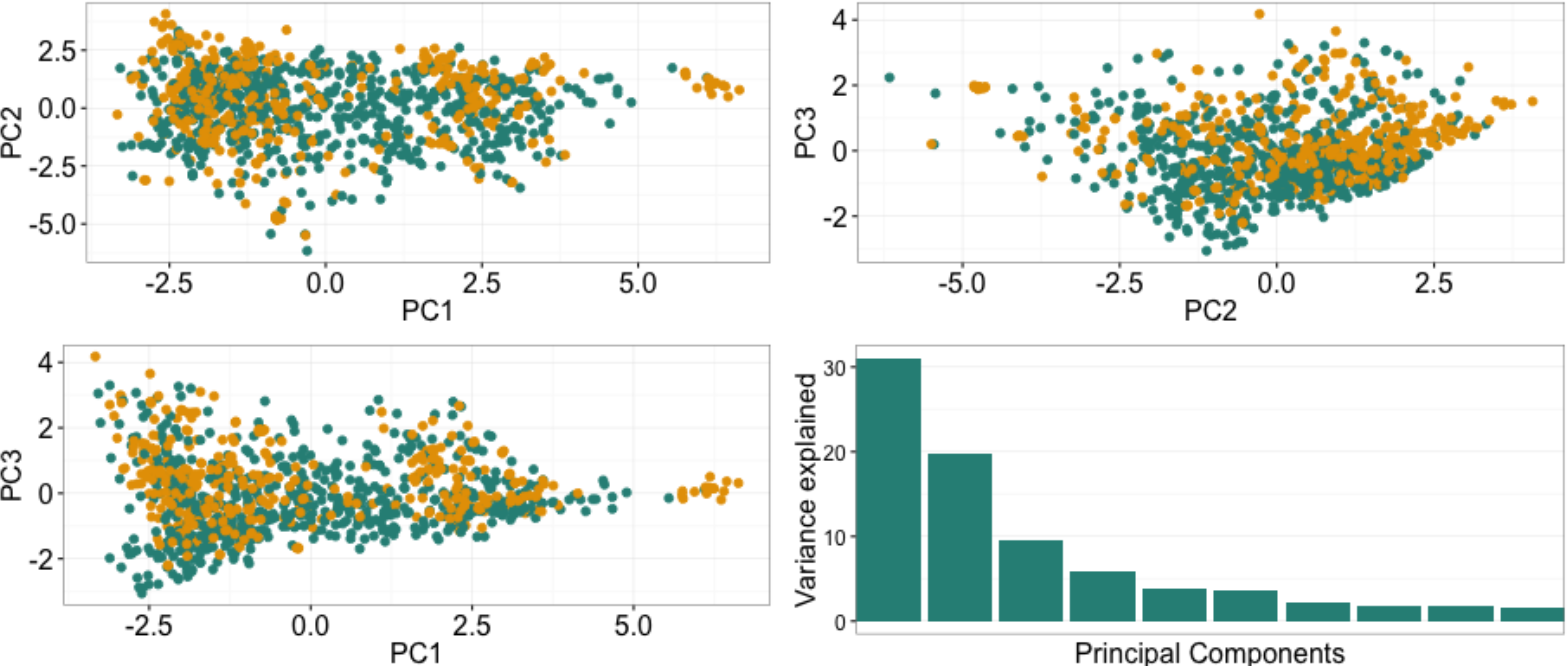
Principal components analysis of panel 1 and panel 2 clones. The top two panels and the lower left panel show the distribution of clones in PC1-PC3. The lower right panel shows the variance explained by the first ten principal components. Green color shows the distribution of panel 1 clones and the orange color shows the distribution of panel 2 clones.

Genotype-phenotype associations for CBSD severity traits based on the combination of multi-location data and 986 individuals are presented in Figure 2. Additional GWAS analyses performed on each panel individually are presented in Supplementary tables 4 and 5 and Supplementary figures 5-12.

**Figure 2.**
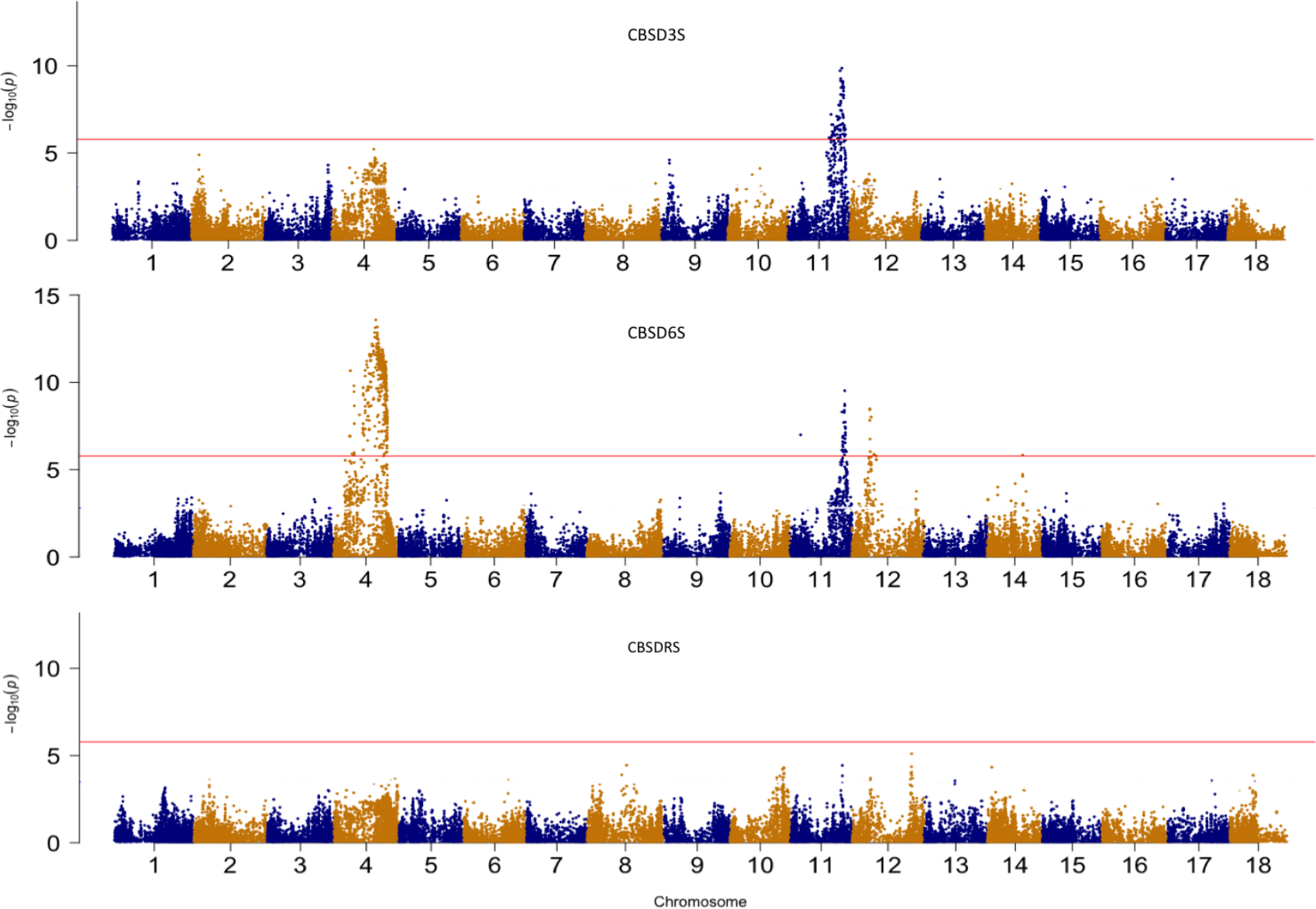
GWAS results for CBSD severity. Analysis was performed with a multilocation combined dataset of panels 1 and 2.(a) scoring 3 MAP (b) 6 MAP and (c) root necrosis severity. Red line indicates Bonferroni threshold.

We characterized SNP markers with a -log10 (P-value) above the Bonferroni threshold > 5.9 as significant marker-trait associations and further annotated those into candidate genes (Supplementary table 4).

For the combined dataset, we identified 83 significant SNP markers associated to CBSD3S; the markers mapped to chromosome 11 with 61 markers located within genes (Supplementary Table 4). The QTL on chromosome 11, top hit reference SNP -log_10_ (P-value) = 9.38, explained 6% of the observed phenotypic variation.

On the other hand, for CBSD6S, we identified significant SNPs on chromosome 11, chromosome 4 and chromosome 12. On chromosome 11, 33 SNPs surpassed the Bonferroni threshold with 27 SNP markers located within genes. The QTL on chromosome 11 is located on the same region as the QTL identified for CBSD3S and explained 5% of the observed phenotypic variation (Figure 3 A).

**Figure 3.**
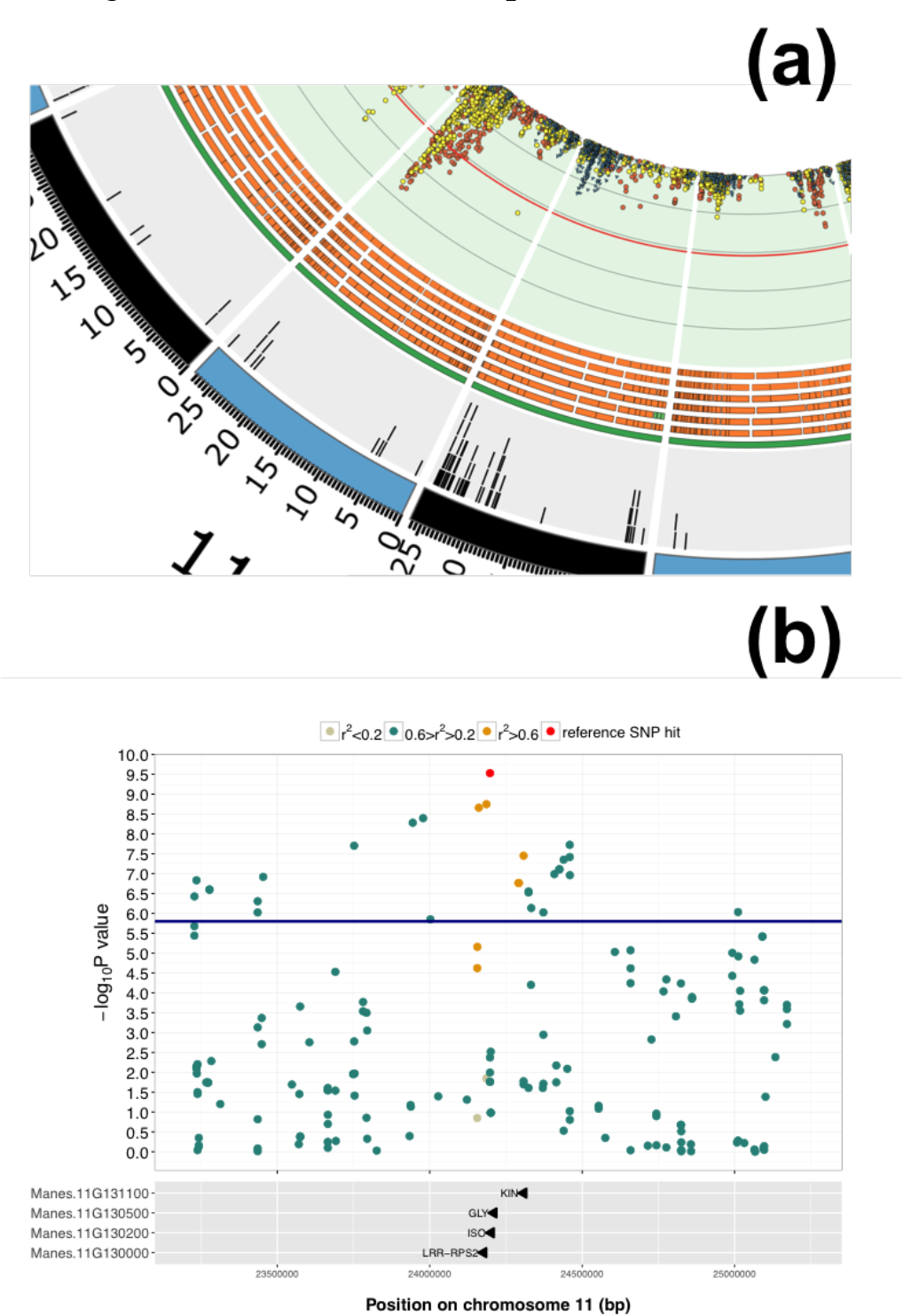
Chromosome 11 region with QTL for CBSD severity. (a) 3 MAP (yellow), 6 MAP and root necrosis (blue).Outer ring black lines indicate clusters of NBS-LRR genes (Lozano et al 2015). Intermediate ring indicate regions homozygous (G/G)(blue) or heterozygous (G/E)(green) for *M. glaziovii* allele and the proportion that were homozygous for the *M. esculenta* allele (E/E)(orange) on seven clones. (b) LD association plot, 2 Mb region in chromosome 11, top SNP indicated in red, annotated genes within that region are indicated in the panel below.

It suffices to note that although several SNPs on chromosome 11 for CBSD6S exceeded the Bonferroni threshold, six SNPs were in linkage disequilibrium (r^2^ > 0.6) with the top reference SNP hit. The SNP markers, with an r^2^> 0.2 to the reference SNP, were annotated into candidate genes: Manes11G130500, a gene that is known to encode glycine-rich protein. Manes11G130000 gene that encodes Leucine-rich repeat (LRR) containing protein, Manes11G130200 gene that encodes the trigger factor chaperone and *peptidyl-prolyl* trans and Manes11G131100 that encodes a protein kinase (Figure 3B).

Since several SNPs on the chromosome 4 QTL region are in high LD, no single locus can be highlighted as candidate gene(s) to be associated with CBSD severity (Figure 4A). The large haplotype on chromosome 4 is an introgression block from the a wild relative of cassava (*M. glaziovii*) (Jennings, 1959; Bredeson *et al.*, 2016). We further confirmed the presence and segregation of the introgressed genome segment in both panels using a set of diagnostic markers from *M. glaziovii* (Figure 4B, supplementary figure 13 and 14).

**Figure 4.**
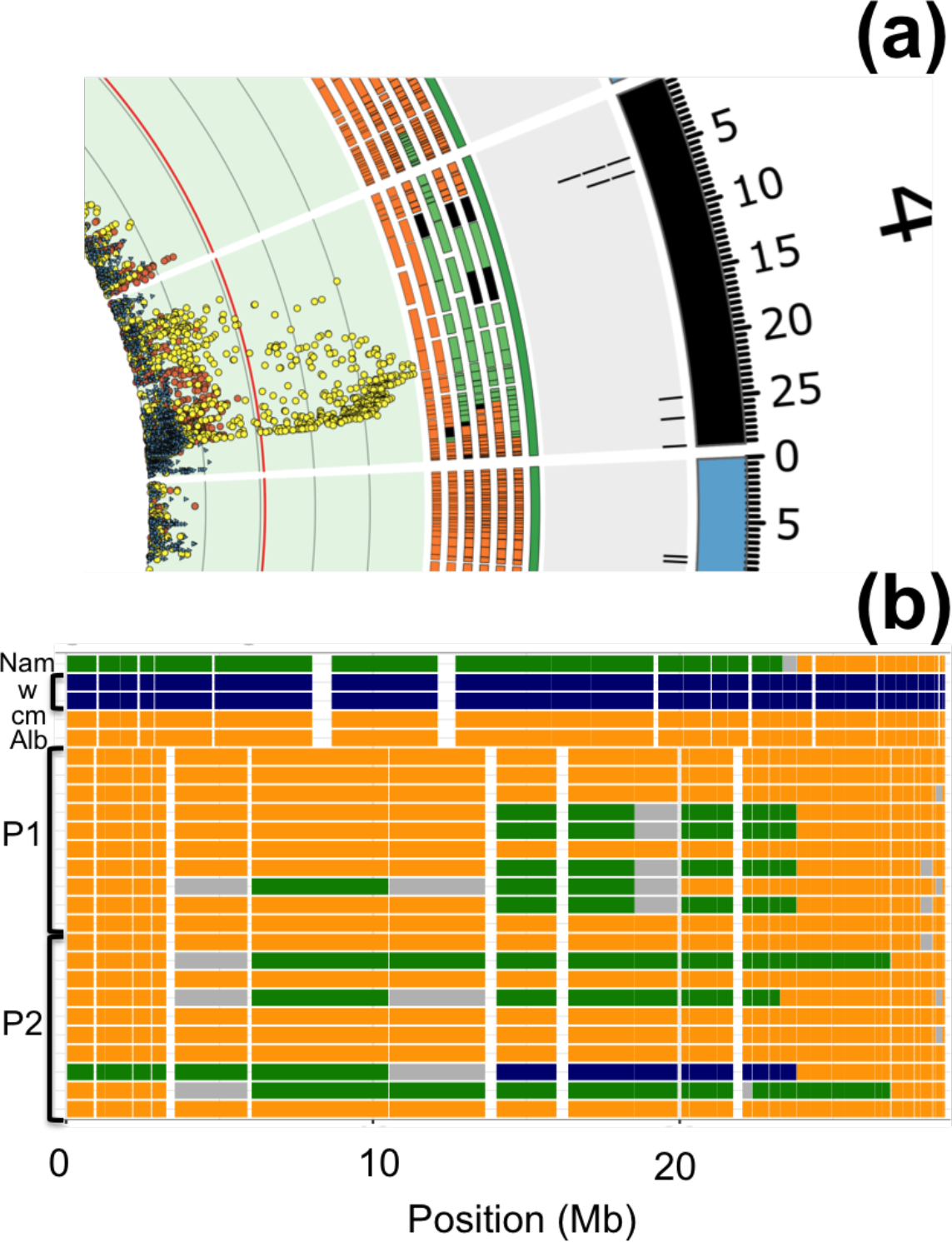
Chromosome 4 region with QTL for CBSD severity with introgression segment. (a) 3 MAP (yellow), 6 MAP and root necrosis (blue). Outer ring black lines indicate clusters of NBS-LRR genes (Lozano et al 2015). Intermediate ring indicate regions homozygous (G/G)(blue) or heterozygous (G/E)(green) for *M. glaziovii* allele and the proportion that were homozygous for the *M. esculenta* allele (E/E)(orange) on seven clones. (b) Introgression region on chromosome 4 (colors description) are the same as the aforementioned),Nam: Namikonga,w: wild *M. glaziovii*, cm: CM330645,Alb:Albert, P1: panel 1 clones and P2 panel 2 clones.

The significant QTL on chromosome 12 has been previously identified for CMD resistance in cassava (Wolfe *et al.*, 2016) Accordingly, after correction for CMD scoring in the first step calculation of CBSD deregressed BLUPs, the QTL on chromosome 12 was no longer significant and only QTLs on chromosomes 4 and 11 remained (supplementary Figure 15).

For CBSDRS we could not identify SNPs surpassing the Bonferroni correction partly to the complexity of this trait with apparently several small effect genes and low heritability. However, the results of the analysis of CBSDRS multi-location data of panel 1 identified significant regions on chromosomes 5, 11 and 18 (-log10 (P-value) > 6.5), which explained 8, 6 and 10% phenotypic variance respectively.

### Genome-wide prediction for CBSV severity in cassava

An important objective within this study was to assess the accuracy of prediction in cassava for CBSD-related traits. Using the combined dataset, we compared the performance of seven genomic prediction models with contrasting assumptions on trait genetic architecture. Some model predictions represent genomic estimated breeding values (GEBV) in that they are sums of additive effects of markers, while other model predictions represent genomic estimated total genetic value (GETGV) because they include non-additive effects. Predictive accuracy for CBSD related traits had mean values across methods of 0.29 (CBSD3S), 0.40 (CBSD6S) and 0.34 (CBSDRS) (Figure 5 and Supplementary table 6).

**Figure 5.**
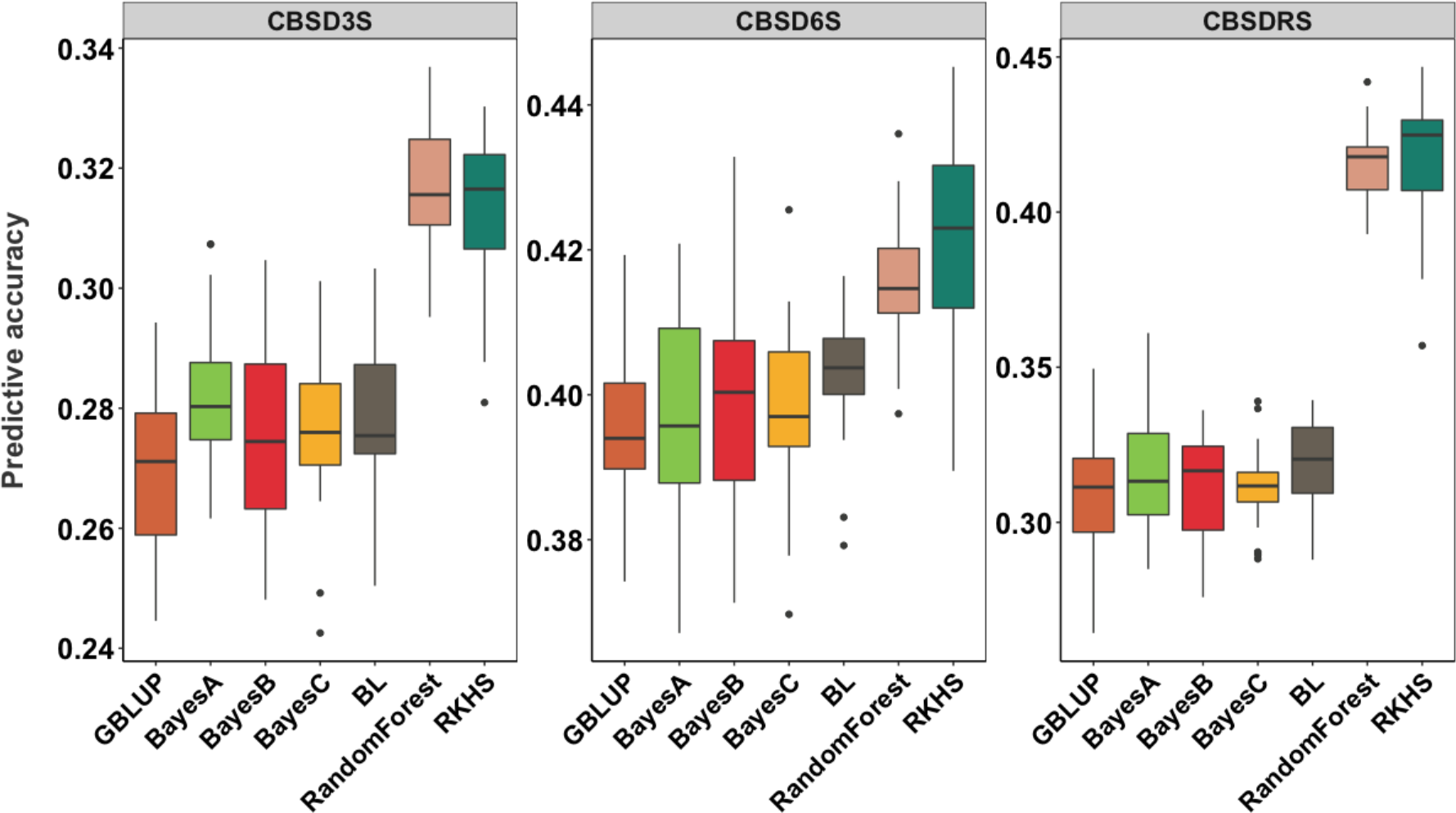
Cross validation results for CBSD severity. 3 MAP (CBSD3S), 6 MAP (CBSD6S) and Root necrosis (CBSDRS). x-axis: predictive accuracy and y-axis: genomic prediction model.

Predictive accuracies for CBSD3S varied in the range of 0.27 (BayesB and GBLUP) and 0.32 (RF), for CBSD6S we obtained a predictive value of 0.40 for most methods except for RKHS (0.42) and RF (0.41) and for CBSD root severity scores varied from 0.31 (BayesA, B, C and GBLUP) to 0.42 (RF and RKHS). It is clear from the results that higher predictive accuracies were consistently achieved when using Random forest and RKHS for the prediction of both foliar and root CBSD resistance traits. Although for foliar symptoms the increase in predictive accuracy using those methods is modest, for CBSDRS the increase in predictive accuracy was 0.10.

Based on the GWAS results, we identified for CBSD3S, CBSD6S and CBSDRS the strongest marker associations on chromosomes 4 and 11. Markers from chromosomes 4, 11 and markers on other chromosomes were used independently to construct covariance matrices that were fitted in a multikernel GBLUP model (Supplementary figure 16). For all CBSD traits the mean predictive accuracy values from the single-kernel GBLUP model were similar to the mean total predictive accuracy following the multi-kernel approach (Supplementary table 6).

Differences were found on the contribution of the individual kernels to the total predictive accuracies. For example, the multikernel GBLUP model for CBSD3S had the lowest total predictive accuracy (0.27) with the highest contribution coming from chromosome 11 and the rest of the genome (0.19). In contrast, the multikernel GBLUP model for CBSD6S gave the highest predictive accuracy (0.40) and most of the accuracy came from chromosome 4 (0.29). The multikernel GBLUP approach for CBSDRS had a total predictive accuracy of 0.30 with the rest of the genome (0.29) contributing the most to the total predictive accuracy (Supplementary figure 16).

## DISCUSSION

Cassava brown streak disease has been identified as one of the most serious threats to food security (Pennisi, 2010) owing to the significant loses it imparts in cassava wherever it occurs. Host plant resistance, that is obtained through breeding efforts has been so far the most effective approach. However, this is only achievable when the host-pathogen behaviour and interaction is well understood and/or when the genetics of resistance to CBSD are clearly known.

In the present study, ∼1200 cassava clones from the NaCRRI breeding program in Uganda were evaluated for CBSD severity scores in leaves and root. Specifically, this paper sought to provide fundamental information on the genetics of resistance to CBSD which was previously unknown. From our analyses it was evident that correlation among foliar CBSD severities were higher than correlation between foliar and root severities.

Selection of resistant clones has been hampered by the fact that some clones do not show symptoms on leaves or storage roots, while other varieties may only express symptoms on leaves and not on roots and still others do not show symptoms on leaves but instead on roots only (ASARECA:, 2013). Moreover, a lack of correlation between virus load and symptom expression in a field evaluation of selected cassava genotypes has been reported (Kaweesi *et al.*, 2014). Previous studies have also reported that 79% plants with above-ground symptoms of CBSD also exhibited root necrosis and 18% of plants had no visible symptoms of CBSV (Hillocks *et al.*, 1996)

Recently, efforts to understand CBSD have focused on CBSD resistance population development and preliminary insights into chromosomal regions and genes involved in resistance (Kawuki *et al.*, 2016; Anjanappa *et al.*, 2016, 2017). These studies have highlighted the existence of a QTL on chromosome 11 for CBSD root necrosis among cassava clones of Tanzanian origin (Kawuki *et al.*, 2016).

In our study, based on foliar CBSD severity scoring using a multi-location dataset we identified significant QTL regions on chromosome 4 and 11, though these associations were not always consistent when the panels were analyzed separately and per location. Overall, these results highlight the advantage of using a large GWAS panel and a multi-location approach were plants are exposed to different disease pressures to identify additional genomic regions.

On chromosome 11, a cluster of genes underlies the significant QTL; candidate genes for further study are: Manes11G131100, Manes11G130500, Manes11G130200 and Manes11G130000. Lozano et al. 2015 previously reported Manes11G130000 when studying the distribution of *NBS-LRR* in cassava. Furthermore, a recent study on early transcriptome response to brown streak virus infection in susceptible and resistant cassava varieties identified Manes.11G130000 among the differentially expressed genes in the susceptible line 60444 from the ETH cassava germplasm collection (Anjanappa *et al.*, 2017). The QTL on chromosome 11 is particularly unstable across locations, which may be related to NBS-LRR genes conferring resistance to a particular strain, UCBSV exhibits a lower mutation rate, while CBSV is more aggressive and mutates faster.

Throughout the 1940s and 1950s at the Amani Research Station, *Manihot glaziovii* and cassava varieties of Brazilian origin were used for crosses to obtain CBSD resistant varieties (Jennings and Iglesias, 2002). One of the introgression segments from these wild relatives has been reported to be located on chromosome 4, however the level of linkage disequilibrium in that region remains as a major constraint for the identification of the gene or genes that are responsible for CBSD resistance (Bredeson *et al.*, 2016). Current on-going research efforts are focused on dissecting the extent of the effects of wild introgressions on cassava traits (Marnin Wolfe personal communication).

One important objective of the present study was to test our ability to predict CBSD severity in cassava, which is, particularly relevant in two situations. First, when the objective is the introduction of germplasm from Latin america and/or from West Africa to East Africa and for early seedling or clonal selection of resistant lines.

Thus, using a cross-validation approach, we evaluated the suitability of seven GS models with the expectation that the results may differ due to differences in genetics of foliar and root CBSD severity traits (B. J. Hayes *et al.*, 2009; Grattapaglia *et al.*, 2011).

In cassava, previous genomic prediction studies have evaluated the predictive ability of GBLUP using historical phenotypic data from the International Institute of Tropical Agriculture (IITA) and GBS markers and in a small training population with relatively low-density markers (de Oliveira *et al.*, 2012; Ly, Hamblin, Rabbi, Melaku, Bakare, Okechukwu, *et al.*, 2013).

Principally, the GS models evaluated have varying underlying assumptions genomic-BLUP (GBLUP) model assume an infinitesimal genetic architecture; Bayesian methods such as BayesA and BayesB relax the assumption of common variance across marker effects (De Los Campos *et al.*, 2009; Habier *et al.*, 2011; Legarra *et al.*, 2014), RKHS and random forest methods can model epistatic and other non-additive effects.

A first assessment of predictive accuracy of CBSD foliar and root traits in cassava indicate that the use of genomic selection is a promising breeding method for resistance to Cassava brown streak virus. We found moderate to high predictive accuracies for these traits in relation to results from other traits in cassava (Ly, Hamblin, Rabbi, Melaku, Bakare, Okechukwu, *et al.*, 2013). However, predictive accuracy values are lower in comparison to the values reported for cassava mosaic virus (Ly, Hamblin, Rabbi, Melaku, Bakare, Okechukwu, *et al.*, 2013) possibly due the presence of a large effect GWAS QTL (CMD2) for CMD.

Although, a priori knowledge of the loci affecting a trait is not needed for GS, we also tested a multiple kernel approach using GWAS results as a reference to construct covariance matrices. GWAS results have been incorporated in genome-wide prediction models to increase predictive accuracy through *de-novo* GWAS or using previously published GWAS results (Zhang *et al.*, 2014; Spindel *et al.*, 2016).

In our study, to avoid a correlation effect across covariance matrices we partitioned SNP markers into three sets: markers on GWAS QTLs chromosomes (chr 4 and 11) and markers on rest of the genome to built genomic relationship matrices (G_chr4_,G_chr11_,G_allchr-[4,11]_). Remarkably, the predictive accuracy of each kernel modeled the genetic architecture found though GWA analyses. Our GWAS and GS results indicate that resistance to CBSD root necrosis severity is polygenic in nature, which is in accordance to Kawuki et al.’s (2016) results.

Our results suggest that non-additive effects are likely to play a role shaping CBSD resistance particularly root necrosis. This conclusion derives from GS results using Random Forest and RKHS, which gave the highest predictive accuracies, and from the observed differences in broad sense and narrow sense heritability values.

CBSD is a disease that has devastating consequences in cassava production and poses a risk particularly to countries in Central and West Africa where CBSD is not currently present. Our study provides, through GWAS and genomic prediction, an insight into the genetic regulation of CBSD severity in leaves and roots. Although we were able to identify a candidate NBS-LRR gene on chromosome 11, the function of this gene in CBSD resistance requires further validation and more importantly, there is a risk that this gene might not be a source of durable resistance to CBSVs. Within this context, genomic selection arises as a promising tool that can accelerate breeding, though the average predictive accuracy is lower than CMD, this is highly variable across locations and the breeding panel evaluated. Further work will require screening of large diversity panels in multiple environments, identification of QTLs specific to viral strains and the introgression of genomic regions conferring resistance to CBSD from wild relatives and Latin American accessions.

## Acknowledgements

We acknowledge the Bill & Melinda Gates Foundation and UKaid (Grant 1048542; http://www.gatesfoundation.org and support from the CGIAR Research Program on Roots, Tubers and Bananas (http://www.rtb.cgiar.org). Thanks also the technical team NaCRRI for collection of phenotypic data

---

**Supplementary figure 1**. Cassava brown streak disease symptoms on leaves and roots of sampled plants; Severity Score from 1 (no visible symptoms) to 5 (severely disease plants. **(a)** leaf veins chlorosis severity progresses with severity score, **(b**) dark brown necrotic areas within storage roots severity scale.

**Supplementary figure 2.** Panel 1 phenotypic distribution of CBSD severity traits. ( A) deregressed BLUPs distribution of CBSD 3 months foliar severity, (B) deregressed BLUPs distribution of CBSD 6 months foliar severity, (C) deregressed BLUPs distribution of CBSD 12 months root severity

**Supplementary figure 3.** Panel 2 phenotypic distribution of CBSD severity traits. (A) deregressed BLUPs distribution of CBSD 3 months foliar severity, (B) deregressed BLUPs distribution of CBSD 6 months foliar severity, (C) deregressed BLUPs distribution of CBSD 9 months foliar severity, (D) deregressed BLUPs distribution of CBSD 12 months root severity

**Supplementary figure 4. Correlation plots between de-regressed BLUPs for foliar and root symptoms**. De-regressed BLUPs were calculated for different locations in panel 1 and panel 2.

**Supplementary figure 5**. GWAS results for CBSD severity in panel 1 measure at Kasese.(a) scoring CBSD 3 months foliar severity (b) 6 CBSD 6 months foliar severity and (c) root necrosis severity. Red line Bonferroni correction. Blue line log_10_ P-value = 3.8.

**Supplementary figure 6**. GWAS results for CBSD severity in panel 1 measure at Ngetta.(a) scoring CBSD 3 months foliar severity (b) 6 CBSD 6 months foliar severity and (c) root necrosis severity. Red line Bonferroni correction. Blue line log_10_ P-value = 3.8.

**Supplementary figure 7**. GWAS results for CBSD severity in panel 1 measure at Namulonge. (a) scoring CBSD 3 months foliar severity (b) 6 CBSD 6 months foliar severity and (c) root necrosis severity. Red line Bonferroni correction. Blue line log_10_ P-value = 3.8.

**Supplementary figure 8**. GWAS results for CBSD severity in with a multilocation dataset of panel 1 (a) scoring CBSD 3 months foliar severity (b) 6 CBSD 6 months foliar severity and (c) root necrosis severity. Red line Bonferroni correction. Blue line logi_0_ P-value = 3.8.

**Supplementary figure 9**. GWAS results for CBSD severity in panel 2 measure at Kamuli. (a) scoring CBSD 3 months foliar severity (b) 6 CBSD 6 months foliar severity (c) 9 CBSD 9 months foliar and (c) root necrosis severity. Red line Bonferroni correction. Blue line log_10_ P-value = 3.8.

**Supplementary figure 10**. GWAS results for CBSD severity in panel 2 measure at Namulonge. (a) Scoring CBSD 3 months foliar severity (b) 6 CBSD 6 months foliar severity (c) 9 CBSD 9 months foliar and (c) root necrosis severity. Red line Bonferroni correction. Blue line log_10_ P-value = 3.8.

**Supplementary figure 11**. GWAS results for CBSD severity in panel 2 at Serere. (a) Scoring CBSD 3 months foliar severity (b) 6 CBSD 6 months foliar severity (c) 9 CBSD 9 months foliar and (c) root necrosis severity. Red line Bonferroni correction. Blue line log_10_ P-value = 3.8.

**Supplementary figure 12**. GWAS results for CBSD severity in with a multilocation dataset of panel 2 (a) scoring CBSD 3 months foliar severity (b) 6 CBSD 6 months foliar severity (c) 9 CBSD 9 months foliar and (c) root necrosis severity. Red line Bonferroni correction. Blue line log_10_ P-value = 3.8.

**Supplementary figure 13.** local LD in chromosome 4. Plot of the mean LD score for each marker. With a smooth line representing a relative measure of the local LD in chromosome 4. Dots are colored with the -log_10_ P-value for the association test for CBSD severity six months after planting.

**Supplementary figure 14.** Introgressions segment detection. For each clone of the two GWAS panels we calculated the proportion of genotypes that were homozygous (G/G) or heterozygous (G/E) for *M. glaziovii* allele and the proportion that were homozygous for the *M. esculenta* allele (E/E).

**Supplementary figure 15. (a)** GWAS results for 6MAP CBSD severity panels 1 and 2 **(b)** GWAS Results after correction including markers in chromosome 12 as a covariate.

**Supplementary figure 16.** Multi-kernel GBLUP approach by fitting three kernels constructed with non-overlapping SNPs (MAF> 0.01) from chromosomes 4, 11 and SNPs from the other chromosomes. Crossvalidation GS predictive accuracies results for CBSD severity were calculated using the multilocation dataset of the combined panels. Scoring CBSD 3 months foliar severity (CBSD3S),CBSD 6 months foliar severity (CBSD6S) and root necrosis severity (CBSDRS).

**Supplementary Table 1**.Pedigree information from GWAS panels 1 and 2. Details are shown on the parental lines per clone and selected traits that came from the maternal side.

**Supplementary table 2**. Correlation values across locations in panel 1 and panel 2. (A) Correlation of deregressed BLUPs across locations within traits in panel 1 measured in three locations.(B) Correlation of deregressed BLUPs across locations within traits in panel 2 measured in three locations

**Supplementary table 3**. Correlation values across locations and traits in panel 1 and panel 2. (A) Correlation of deregressed BLUPs across locations and traits in panel 1 measured in three locations.(B) Correlation of deregressed BLUPs across locations and four traits in panel 2 measured in three locations

**Supplementary table 4**. Panel 1 and 2 and combined panels GWAS results. Gene annotation is only shown for significant SNPs.

**Supplementary table 5**. Explained variance of phenotypic traits. Details are shown of the reference SNP, the -log10(pval)(score),chromosome and explained variance.

**Supplementary table 6**.Genomic prediction accuracy values. (A) Cross validation results using 7 GS models for CBSD severity prediction of 3 MAP CBSD3S, 6 MAP CBSD6S and Root necrosis (CBSDRS) (B) Multi-kernel GBLUP crossvalidation by fitting three kernels constructed with non-overlapping SNPs (MAF> 0.01) from chromosomes 4, 11 and SNPs from the other chromosomes. RKHS = Reproducing kernel Hilbert spaces regression, Total accuracy is the accuracy obtained by following the GBLUP multikernel approach.

## References

Akdemir D, Okeke UG (2015). EMMREML: Fitting Mixed Models with Known Covariance Structures.

Alicai T, Ndunguru J, Sseruwagi P, Tairo F, Okao-Okuja G, Nanvubya R, et al. (2016). Characterization by Next Generation Sequencing Reveals the Molecular Mechanisms Driving the Faster Evolutionary rate of Cassava brown streak virus Compared with Ugandan cassava brown streak virus. Cold Spring Harbor Labs Journals.

Anjanappa RB, Mehta D, Maruthi MN, Kanju E, Gruissem W, Vanderschuren H (2016). Characterization of Brown Streak Virus-Resistant Cassava. 29: 527–534.

Anjanappa RB, Mehta D, Okoniewski MJ, Szabelska A, Gruissem W, Vanderschuren H (2017). Early transcriptome response to brown streak virus infection in susceptible and resistant cassava varieties.: 1–22.

ASARECA: (2013). ASARECA Annual Report 2012: Transforming Agriculture for Economic Growth in Eastern and Central Africa. Entebbe, Uganda.

B. J. Hayes, H. D. Daetwyler, P. Bowman, G. Moser, B. Tier4, R. Crump, et al. (2009). Accuracy of Genomic Selection: Comparing Theory and Results. In: Daetwyler HD (ed) Genome-Wide Evaluation of Populations, Proc. of Assoc. Advmt. Anim. Breed.,pp 352–355.

Bredeson J V, Lyons JB, Prochnik SE, Wu GA, Ha CM, Edsinger-Gonzales E, et al. (2016). Sequencing wild and cultivated cassava and related species reveals extensive interspecific hybridization and genetic diversity. Nat Biotechnol 34: 562–570.

Breiman L (2001). Random Forests. Mach Learn 45: 5–32.

Browning BL, Browning SR(2016). Genotype Imputation with Millions of Reference Samples. Am J Hum Genet 98: 116–126.

Bulik-Sullivan B, Finucane HK, Anttila V, Gusev A, Day FR, Loh P-R, et al. (2015). An atlas of genetic correlations across human diseases and traits. Nat Publ Gr 47.

Charmet G, Storlie E (2012). Implementation of genome-wide selection in wheat. Russ J Genet Appl Res 2: 298–303.

Charmet G, Storlie E, Oury FX, Laurent V, Beghin D, Chevarin L, et al. (2014).Genome-wide prediction of three important traits in bread wheat. Mol Breed 34: 1843–1852.

Cros D, Denis M, Sánchez L, Cochard B, Flori A, Durand-gasselin T, et al. (2015). Genomic selection prediction accuracy in a perennial crop: case study of oil palm (Elaeis guineensis Jacq.). Theor Appl Genet 128: 397–410.

Elshire RJ, Glaubitz JC, Sun Q, Poland J a, Kawamoto K, Buckler ES, et al. (2011). A robust, simple genotyping-by-sequencing (GBS) approach for high diversity species. PLoS One 6: e19379.

Endelman JB (2011). Ridge Regression and Other Kernels for Genomic Selection with R Package rrBLUP. Plant Genome J 4: 250.

Esuma W, Herselman L, Labuschagne MT, Ramu P, Lu F, Baguma Y, et al. (2016). Genome-wide association mapping of provitamin A carotenoid content in cassava. Euphytica.

Federer WT, Crossa J (2012). Screening Experimental Designs for Quantitative Trait Loci, Association Mapping, Genotype-by Environment Interaction, and Other Investigations. Front Physiol 3.

Federer WT, Nguyen N-K, others (2002). Constructing Augmented Experiment Designs with Gendex.

Garrick DJ, Taylor JF, Fernando RL (2009). Deregressing estimated breeding values and weighting information for genomic regression analyses. Genet Sel Evol 41: 55.

Glaubitz JC, Casstevens TM, Lu F, Harriman J, Elshire RJ, Sun Q, et al. (2014). TASSEL-GBS: a high capacity genotyping by sequencing analysis pipeline. PLoS One 9: e90346.

Goodstein D, Batra S, Carlson J, Hayes R, Phillips J, Shu S, et al. (2014). Phytozome Comparative Plant Genomics Portal.

Gouy M, Rousselle Y, Bastianelli D, Lecomte P, Bonnal o L, Roques D, et al. (2013). Experimental assessment of the accuracy of genomic selection in sugarcane. Theor Appl Genet 126: 1–12.

Grattapaglia D, Deon M, Resende V, Resende MR, Sansaloni CP, Petroli CD, et al. (2011). Genomic Selection for growth traits in Eucalyptus: accuracy within and across breeding populations. From IUFRO Tree Biotechnol Conf BMC Proc 5.

Habier D, Fernando RL, Kizilkaya K, Garrick DJ (2011). Extension of the bayesian alphabet for genomic selection. BMC Bioinformatics 12: 186.

Hamblin MT, Rabbi IY (2014). The Effects of Restriction-Enzyme Choice on Properties of Genotyping-by-Sequencing Libraries: A Study in Cassava (). Crop Sci 0: 0.

Heslot N, Akdemir D, Sorrells ME, Jannink JL (2014). Integrating environmental covariates and crop modeling into the genomic selection framework to predict genotype by environment interactions. Theor Appl Genet 127: 463–480.

Heslot N, Yang H-P, Sorrells ME, Jannink J-L (2012). Genomic Selection in Plant Breeding: A Comparison of Models. Crop Sci 52: 146.

Hillocks RJ, Jennings DL (2003). Cassava brown streak disease: a review of present knowledge and research needs. Int J Pest Manag 49: 225–234.

Hillocks RJ, Raya M, Thresh JM (1996). The association between root necrosis and above-ground symptoms of brown streak virus infection of cassava in southern Tanzania. Int J Pest Manag 42: 285–289.

Jannink J-LL, Lorenz AJ, Iwata H (2010). Genomic selection in plant breeding: from theory to practice. Brief Funct Genomics 9: 166–177.

Jennings DL (1959). Manihot melanobasis Müll. Arg.—a useful parent for cassava breeding. Euphytica 8: 157–162.

Jennings DL, Iglesias C (2002). Breeding for Crop Improvement. Cassava Biol Prod Util: 149–166.

Kaweesi T, Kawuki R, Kyaligonza V, Baguma Y, Tusiime G, Ferguson ME (2014). Field evaluation of selected cassava genotypes for cassava brown streak disease based on symptom expression and virus load. Virol J 11: 216.

Kawuki RSRS, Kaweesi T, Esuma W, Pariyo A, Kayondo IS, Ozimati A, et al. (2016). Eleven years of breeding efforts to combat cassava brown streak disease. Breed Sci 66: 560–571.

Kulembeka HP (2010). Genetic linkage mapping of Field Resistance to cassava brown streak Disease in cassava landraces from Tanzania. University of the Free State.

Legarra A, Christensen OF, Aguilar I, Misztal I (2014). Single Step, a general approach for genomic selection. Livest Sci 166: 54–65.

Legg J, Somado EA, Barker I, Beach L, Ceballos H, Cuellar W, et al. (2014). A global alliance declaring war on cassava viruses in Africa. Food Secur 6: 231–248.

Legg JP, Sseruwagi P, Boniface S, Okao-Okuja G, Shirima R, Bigirimana S, et al. (2014). Spatiotemporal patterns of genetic change amongst populations of cassava Bemisia tabaci whiteflies driving virus pandemics in East and Central Africa. Virus Res 186: 61–75.

Liaw a, Wiener M (2002). Classification and Regression by random Forest. R news 2: 18–22.

Lorenz AJ, Chao S, Asoro FG, Heffner EL, Hayashi T, Iwata H, et al. (2011). Genomic Selection in Plant Breeding. Knowledge and Prospects. In: Advances in Agronomy, Elsevier Inc Vol 110, pp 77–123.

De Los Campos G, Naya H, Gianola D, Crossa J, Legarra A, Manfredi E, et al. (2009). Predicting quantitative traits with regression models for dense molecular markers and pedigree. Genetics 182: 375–385.

Lozano R, Hamblin MT, Prochnik S, Jannink J-L (2015). Identification and distribution of the NBS-LRR gene family in the Cassava genome. BMC Genomics 16: 1–14.

Ly D, Hamblin M, Rabbi I, Melaku G, Bakare M, Gauch HG, et al. (2013). Relatedness and genotype ?? environment interaction affect prediction accuracies in genomic selection: A study in cassava. Crop Sci 53:1312–1325.

Ly D, Hamblin M, Rabbi I, Melaku G, Bakare M, Okechukwu R, et al. (2013). Relatedness and Genotype-by-environment Interaction Affect Prediction Accuracies in Genomic Selection: a Study in Cassava 2. Crop Sci.

Maruthi MN, Jeremiah CS, Mohammed IU, Legg JP (2016). Virus-vector relationships and the role of whiteflies, Bemisia tabaci, and farmer practices in the spread of cassava brown streak viruses.

Mbanzibwa DR, Tian YP, Tugume AK, Mukasa SB, Tairo F, Kyamanywa S, et al. (2011). Simultaneous virus-specific detection of the two cassava brown streak-associated viruses by RT-PCR reveals wide distribution in East Africa, mixed infections, and infections in Manihot glaziovii. J Virol Methods 171: 394–400.

McQuaid CF, Sseruwagi P, Pariyo A, van den Bosch F (2016). Cassava brown streak disease and the sustainability of a clean seed system. Plant Pathol 65: 299–309.

Meuwissen THE, Hayes BJ, Goddard ME (2001). Prediction of total genetic value using genome-wide dense marker maps. Genetics 157: 1819–1829.

Motsinger-Reif AA, Reif DM, Fanelli TJ, Ritchie MD (2008). A comparison of analytical methods for genetic association studies. Genet Epidemiol 32: 767–778.

Munga TL (2008). Breeding for Cassava Brown Streak Resistance in Coastal Kenya. University of KwaZulu-Natal Republic of South Africa.

Ndunguru J, Sseruwagi P, Tairo F, Stomeo F, Maina S, Djinkeng A, et al. (2015). Analyses of twelve new whole genome sequences of cassava brown streak viruses and ugandan cassava brown streak viruses from East Africa: Diversity, supercomputing and evidence for further speciation. PLoS One 10: e0139321.

Ndyetabula IL, Merumba SM, Jeremiah SC, Kasele S, Mkamilo GS, Kagimbo FM, et al. (2016). Analysis of Interactions Between Cassava Brown Streak Disease Symptom Types Facilitates the Determination of Varietal Responses and Yield Losses. Plant Dis 100: 1388–1396.

Nichols RFW (1947). Breeding cassava for virus resistance. East African Agric J 12: 184–94.

Ogwok E, Patil BL, Alicai T, Fauquet CM (2010). Transmission studies with Cassava brown streak Uganda virus (Potyviridae: Ipomovirus) and its interaction with abiotic and biotic factors in Nicotiana benthamiana. J Virol Methods 169: 296–304.

de Oliveira EJ, de Resende MDV, da Silva Santos V, Ferreira CF, Oliveira GAF, da Silva MS, et al. (2012). Genome-wide selection in cassava. Euphytica 187: 263–276.

Park T, Casella G (2008). The Bayesian Lasso. J Am Stat Assoc 103: 681–686.

Patil BL, Legg JP, Kanju E, Fauquet CM (2015). Cassava brown streak disease: A threat to food security in Africa. J Gen Virol 96: 956–968.

Pennisi E (2010). Armed and dangerous. Science 327: 804–5.

Pérez JC, Lenis JI, Calle F, Morante N, Sánchez T, Debouck D, et al. (2011). Genetic variability of root peel thickness and its influence in extractable starch from cassava (Manihot esculenta Crantz) roots. Plant Breed 130: 688–693.

Pérez P, De Los Campos G (2014). Genome-wide regression and prediction with the BGLR statistical package. Genetics 198: 483–495.

Prochnik S, Marri PR, Desany B, Rabinowicz PD, Kodira C, Mohiuddin M, et al. (2012). The Cassava Genome: Current Progress, Future Directions. Trop Plant Biol 5: 88–94.

Purcell S, Neale B, Todd-Brown K, Thomas L, Ferreira MAR, Bender D, et al. (2007). PLINK: A Tool Set for Whole-Genome Association and Population-Based Linkage Analyses. Am J Hum Genet 81: 559–575.

QIAGEN (2012). DNeasy ^®^ Plant Handbook DNeasy Plant Mini Kit and tissues, or fungi Sample & Assay Technologies QIAGEN Sample and Assay Technologies.

Quinlan AR, Hall IM (2010). BEDTools: a flexible suite of utilities for comparing genomic features. Bioinforma ApplNOTE 26: 841–84210.

Ramu P, Esuma W, Kawuki R, Rabbi IY, Egesi C, Bredeson J V, et al. (2016). Cassava HapMap: Managing genetic load in a clonal crop species. bioRxiv: 1–15.

Rentería ME, Cortes A, Medland SE (2013). Using PLINK for Genome-Wide Association Studies (GWAS) and Data Analysis. In:Gondro C, van der Werf J, Hayes B (eds) Humana Press: Totowa, NJ Vol 1019, pp 193–213.

Resende MFR, Muñoz P, Acosta JJ, Peter GF, Davis JM, Grattapaglia D, et al. (2012). Accelerating the domestication of trees using genomic selection: accuracy of prediction models across ages and environments. New Phytol 193: 617–624.

Spindel JE, Begum H, Akdemir D, Collard B, Redoña E, Jannink J, et al. (2016). Genome-wide prediction models that incorporate de novo GWAS are a powerful new tool for tropical rice improvement. Heredity (Edinb) 116: 395–408.

Spindel J, Begum H, Akdemir D, Virk P, Collard B, Redoña E, et al. (2015). Genomic Selection and Association Mapping in Rice (Oryza sativa): Effect of Trait Genetic Architecture, Training Population Composition, Marker Number and Statistical Model on Accuracy of Rice Genomic Selection in Elite, Tropical Rice Breeding Lines. PLoS Genet 11: 1–25.

Strobl C, Malley J, Tutz G (2009). An Introduction to Recursive Partitioning: Rationale, Application, and Characteristics of Classification and Regression Trees, Bagging, and Random Forests. Psychol Methods 14: 323–348.

Turner SD (2014). qqman: an R package for visualizing GWAS results using Q-Q and manhattan plots.

VanRaden PM (2008). Efficient methods to compute genomic predictions. JDairy Sci 91: 4414–23.

Winter S, Koerbler M, Stein B, Pietruszka A, Paape M, Butgereitt A (2010). Analysis of cassava brown streak viruses reveals the presence of distinct virus species causing cassava brown streak disease in East Africa. J Gen Virol 91: 1365–1372.

Wolfe MD, Rabbi IY, Egesi C, Hamblin M, Kawuki R, Kulakow P, et al. (2016). Genome-wide association and prediction reveals the genetic architecture of cassava mosaic disease resistance and prospects for rapid genetic improvement. Plant Genome 9: 1–13.

Yang J, Lee SH, Goddard ME, Visscher PM (2011). GCTA: A tool for genome-wide complex trait analysis. Am J Hum Genet 88: 76–82.

Zhang Z, Ober U, Erbe M, Zhang H, Gao N, He J, et al. (2014). Improving the accuracy of whole genome prediction for complex traits using the results of genome wide association studies (X, Cai, Ed.). PLoS One 9: e93017.

